# Organismal responses to artificial light at night in terrestrial ecosystems vary across lunar cycles

**DOI:** 10.64898/2026.06.15.732439

**Authors:** John F. Deitsch, Brett M. Seymoure

**Affiliations:** Department of Biological Sciences, The University of Texas at El Paso, El Paso, TX, USA, 79902

## Abstract

Globally, nocturnal lightscapes are now determined by both moonlight and light pollution, or artificial light at night (ALAN). Organisms respond both to changes in moonlight across lunar cycles and to alterations in light conditions due to artificial light. The interaction of natural and artificial light is a critical aspect to incorporate into our understanding of how crepuscular and nocturnal ecology is altered in anthropogenically-modified landscapes. In this manuscript we review the rapidly expanding body of research on ecological impacts of ALAN to (1) assess patterns of lunar data inclusion and (2) summarize documented interactions of moonlight and ALAN. Three-fourths (72%) of 379 papers reviewed did not incorporate moonlight into their statistical analyses and experimental design. Only 12% directly investigated interaction effects of moonlight and ALAN. However, 70% of these studies reported an interactive effect. Considering this stark contrast, as a precursor to our literature review, we present an overview of moonlight and the lunar cycle for biologists. The overarching trend emerging from the literature is that biological impacts of ALAN decrease with increasing moonlight, although the opposite is true in some cases. After summarizing the literature, we present general hypotheses regarding the interaction of the lunar cycle and ALAN. These hypotheses consider the different forms of ALAN encountered by organisms (i.e. skyglow and light sources) and account for the influence of cloud cover. Finally, we suggest best practices for incorporating moonlight into future research on biological impacts of ALAN.

## Introduction

Disruption of natural light cycles (daily light-dark, circa-monthly lunar cycles) by artificial light pollution is a major component of global environmental change (Davies and Smyth 2018). Nightly and monthly variation in moonlight determines natural nighttime light conditions; consequently, light pollution affects organismal behavior and ecosystem functioning (Puschnig et al. 2020, Dominoni et al. 2020). Understanding how organisms respond to the interaction of artificial light and moonlight is critical for contextualizing light pollution’s ecological impacts. Research on ecological impacts of light pollution, or artificial light at night (ALAN), has increased in the past two decades (Davies and Smyth 2018) but the ecological context of moonlight is absent from many of these studies. This manuscript has four primary objectives. First, to outline ecologically relevant aspects of moonlight and the lunar cycle. Second, to assess patterns of lunar inclusion in published research on ALAN’s ecological impacts. Third, to summarize documented interactions of ALAN and moonlight. Fourth, to present a broad hypothetico-deductive framework for future research on ALAN and ecology. In this manuscript, we discuss physical characteristics of moonlight and their variation across lunar cycles to the degree relevant to biology, behavior, and photosystems of organisms. Although important, we do not focus on underlying physics or applications to remote sensing.

## 1 Artificial light at night

Over the past three decades, a substantial body of literature on biological and ecological impacts of artificial light at night (ALAN) has been generated (Davies and Smyth 2018). This research has shown that ALAN’s influence extends across levels of biological organization, from physiology to populations (Sanders et al. 2021). Excellent reviews have highlighted many aspects of ALAN’s biological footprint, including the impact on particular biomes, taxonomic groups, or behaviors (Owens and Lewis 2018, Desouhant et al. 2019, Secondi et al. 2020, Owens et al. 2020, Boyes et al. 2020, Sanders et al. 2021, Miller and Rice 2023, Burt et al. 2023). One element of this facet of global environmental change meriting greater attention is the interaction of ALAN with the primary determinant of natural nighttime light, the lunar cycle.

ALAN is a globally pervasive form of environmental change that is expanding in extent and intensity (Falchi 2016, Kyba et al. 2017a, 2023). In particularly light-polluted locations, ALAN significantly reduces or eliminates circa-monthly lunar cycles in nighttime brightness (Puschnig et al. 2020, Seymoure et al. 2025). However, even comparatively weak ALAN disrupts natural light conditions. Skyglow levels twice as bright as natural new moon conditions – critical conditions for many biological processes – occurs across a quarter of terrestrial land area, including 77% of Globally Protected Areas and 51% of Key Biodiversity Areas (Seymoure et al. 2025).

ALAN is present in the landscape in different forms. In terrestrial biology, ALAN is often investigated as individual light sources, skyglow, or ambient light levels. Individual light sources are precise locations of elevated radiance on the landscape. Skyglow is the artificial brightening of the night sky caused by backscattering of photons emitted by ground-level light sources (Falchi 2016). Individual light sources and downwelling light from skyglow contribute to elevated ambient light levels. Skyglow can lead to increased ambient light levels in locations where individual light sources are not directly visible (Dyer et al. 2023). Organisms respond differently to skyglow and point sources (Foster et al. 2021, Dickerson et al. 2022, Grenis et al. 2023, Barrientos et al. 2023, Hirt et al. 2023). Though the physics of light complicate a rigid classification of ALAN, the disparity in organismal response to different scales of ALAN makes a categorization of ALAN useful for developing research questions and hypotheses (Kehoe et al. 2022, Hirt et al. 2023). One framework is a distinction between ALAN as radiance (light emitted by a light source or a specific point of artificially brightened sky) and ALAN as irradiance (elevated ambient light levels). This distinction is helpful when considering the influence of moonlight on the impacts of ALAN.

Organisms encounter ALAN under markedly different sensory environments, within and across nights, as moonlight changes with lunar phase and elevation angle. To reach a more complete understanding of ALAN’s biological footprint, incorporating lunar information is crucial. Moonlight is often mischaracterized or over-simplified in biological literature (Kyba et al. 2017b, 2020). In the following section, before reporting the results of our literature review, we present an overview of moonlight as an ecological variable.

## 2 Moonlight

### 2.1 Overview

Moonlight is sunlight that has reflected off the lunar surface and reached Earth. Each photon of moonlight has three fundamental characteristics: wavelength, frequency, and polarization. Wavelength and frequency, inversely proportional, determine the energy of a photon (in the visible spectrum, wavelength/frequency determines color perception). Polarization describes the direction of the photon’s oscillating electromagnetic field. Moonlight, of course, is not one photon but a stream of billions of photons. This stream of photons has its own characteristics including intensity, hue, saturation, and polarity (for an excellent review of these principles of light, see Montgomerie 2006). Moonlight can be measured as the amount of light leaving the moon (radiance or luminance) or as the amount of light striking a specified area of the Earth (irradiance or illuminance) (Supplementary Materials 1) (Johnsen 2012). The characteristics of moonlight change drastically due to both the geometry of celestial bodies and environmental variables within Earth’s atmosphere (e.g. cloud cover).

The lunar phase is the primary source of variability in moonlight on earth. Lunar phases arise from the constantly changing geometry amongst the Sun, Earth, and Moon as the Moon orbits the Earth and the Earth orbits the Sun (Figure 1A). The portion of the Moon that is illuminated by the Sun and facing Earth changes as the Moon moves along its orbital path (Figure 1B). When the Moon is ‘between’ the Earth and the Sun, its sunlit side faces away from earth, producing a new moon. When the Earth is ‘between’ the Sun and Moon, the sunlit side faces earth, creating a full moon. Intermediate phases (crescent, quarter, and gibbous) occur as the angle formed by the Moon, Earth, and Sun increases, respectively. The nocturnal ambient light environment (irradiance) on Earth’s surface differs immensely depending upon the lunar phase: full moon nights (up to 300 mlux) can be up to thousand times brighter than new moon nights (0.1–1 mlux), respectively (Duriscoe 2016, Śmielak 2023). However, this 300 mlux values represents only the brightest moonlit nights; 50 to 200 mlux is more typical for full moon conditions (Kyba et al. 2017).

**Figure 1:**
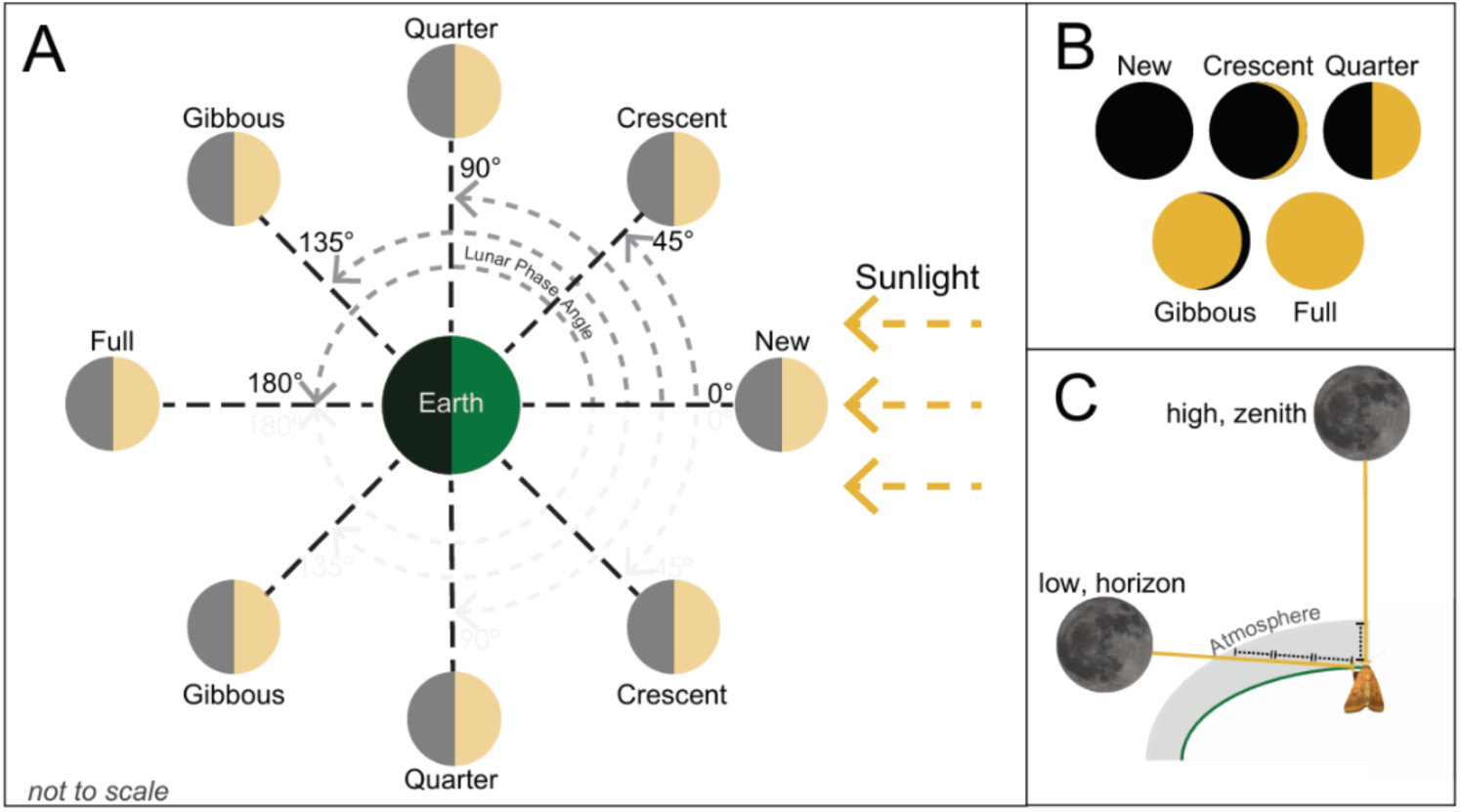
**A)** Celestial positioning of the lunar cycle with lunar phase angle. **B)** Appearance of lunar phases from earth. **C)** Extinction of moonlight in the atmosphere depends on elevation above horizon (perspective of viewer) due to different distances traveled through atmosphere.

Lunar elevation angle, the angular height of the moon above the horizon, is the next largest contributor to variation in moonlight. When the moon is high in the night sky, nocturnal irradiance can be up to 200 times brighter than when the moon is near the horizon (Seymoure et al. 2025). Furthermore, the coloration of moonlight can change with lunar elevation angles. When the Moon is high in the sky, it can appear as a rusty-cream color, akin to amber lighting. However, due to Rayleigh scattering (the scattering of shorter wavelengths of light) this lighting becomes dominated by longer wavelengths and thus reddens as the Moon approaches the horizon (Johnsen 2012).

Light emitted from the sun is unpolarized but becomes partially polarized when scattered by the lunar surface (Venkatesulu and Shaw 2024). This partially polarized moonlight is linearly polarized by scattering while traveling through the earth’s upper atmosphere (Freas and Cheng 2025). This produces a polarization pattern in the night sky that is similar but fainter than that of sunlight (Cronin et al. 2014). Organisms are capable of detecting and utilizing these polarization signals (Muheim et al. 2006, Johnsen 2012, Freas et al. 2017, Foster et al. 2019).

Across the circa-monthly lunar cycle, there is dramatic variation in the magnitude and timing of moonlight. The proportion of nighttime in which the moon is visible above the horizon varies across the lunar cycle (Figure 2A): full moons are visible throughout nighttime while new moons are visible for none of the night. Average nighttime lunar elevation and irradiance are greater during full moons than new moons (Figure 2B, 2C). Two nights with similar lunar irradiance can have markedly different timing of visibility – first quarter and third quarter moons are visible for the first and second half of nighttime, respectively (Figure 2D).

**Figure 2:**
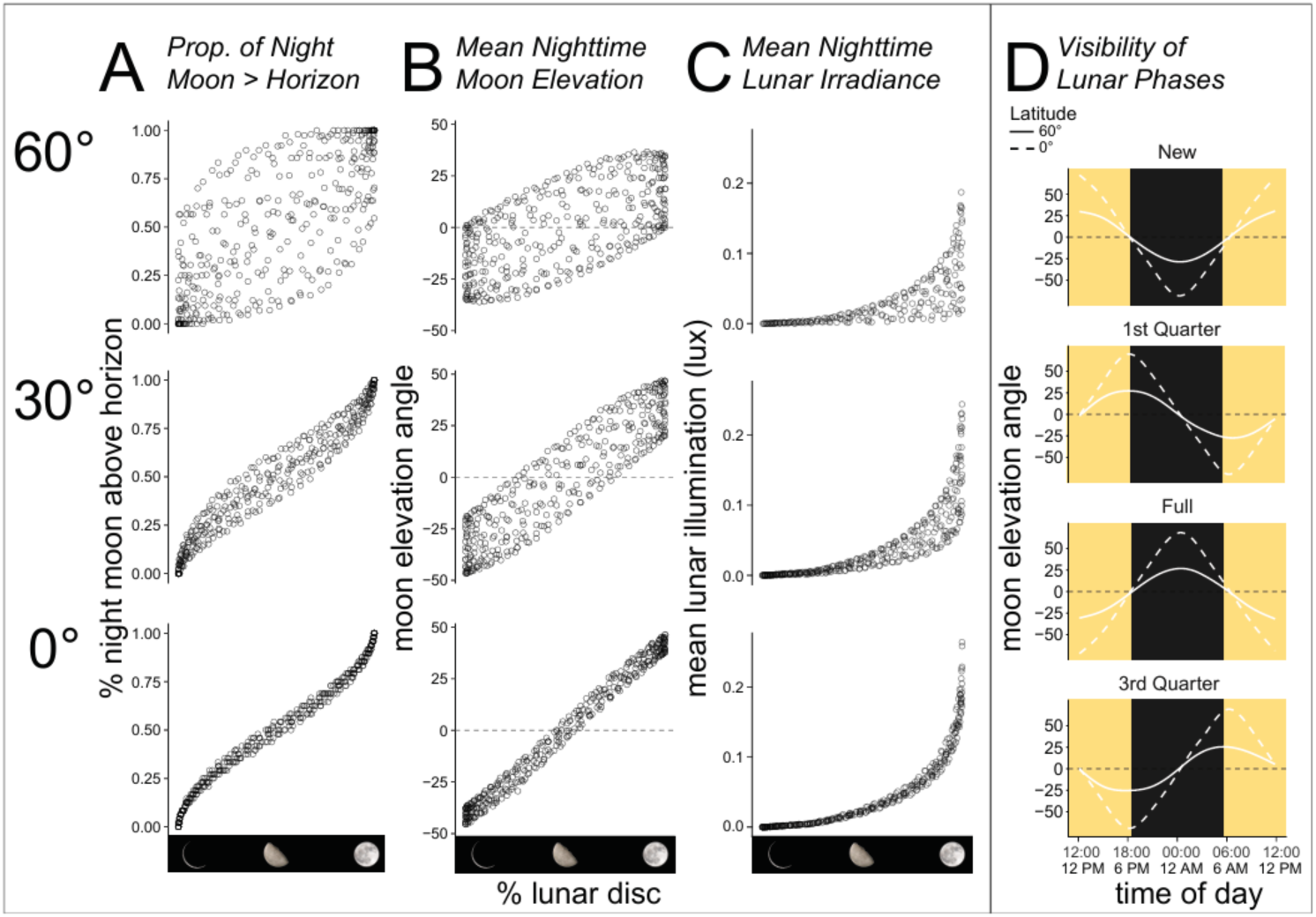
Panels A, B, and C represent one year (2025) of nightly data generated via the *moonlit* R package for 3 latitudes. Each point represents one night. **A)** Proportion of night that moon is visible above horizon. **B)** Average nighttime moon elevation angle above horizon. **C)** Average modeled nighttime lunar irradiance (lux). **D)** Timing of visibility of moon above horizon for four primary lunar phases (new, full, first quarter, third quarter) at two latitudes. The x-axis is centered at midnight (0:00).

### 2.2 Properties of Moonlight

#### 2.2.1 Lunar Radiance

Lunar radiance is the amount of sunlight reflected by the lunar surface. Colloquially, this is often referred to as the “brightness” of the moon. Lunar radiance depends on lunar albedo: the fraction of incident sunlight that is reflected by the lunar surface (Lane and Irvine 1973). Lunar albedo increases as the lunar phase trends towards a full moon, with a rapid increase immediately before a full moon due to the opposition effect/surge, i.e. the increase in albedo as a result of coherent backscattering and shadow hiding (Kieffer and Stone 2005, Śmielak 2023). The relationship between lunar radiance and lunar phase is non-linear.

#### 2.2.2 Lunar Irradiance

Lunar irradiance is the amount of reflected sunlight received by a location on earth. Lunar irradiance depends on the fraction of the lunar disc reflecting sunlight and the elevation angle of the moon above the horizon (Figure 3). Lunar irradiance increases with lunar phase and with elevation; a zenith full moon provides maximal lunar irradiance in a landscape. When the moon is close to the horizon, moonlight travels a greater distance through the earth’s atmosphere (Figure 1C) before reaching the observer and is subject to increased scattering and extinction (Śmielak 2023). Since both lunar irradiance and the proportion of nighttime with the moon over the horizon increase with lunar phase, the total moonlight received by an ecosystem is drastically higher on a full moon night. Cloud cover reduces lunar irradiance at ground level (Kyba et al. 2011a).

**Figure 3:**
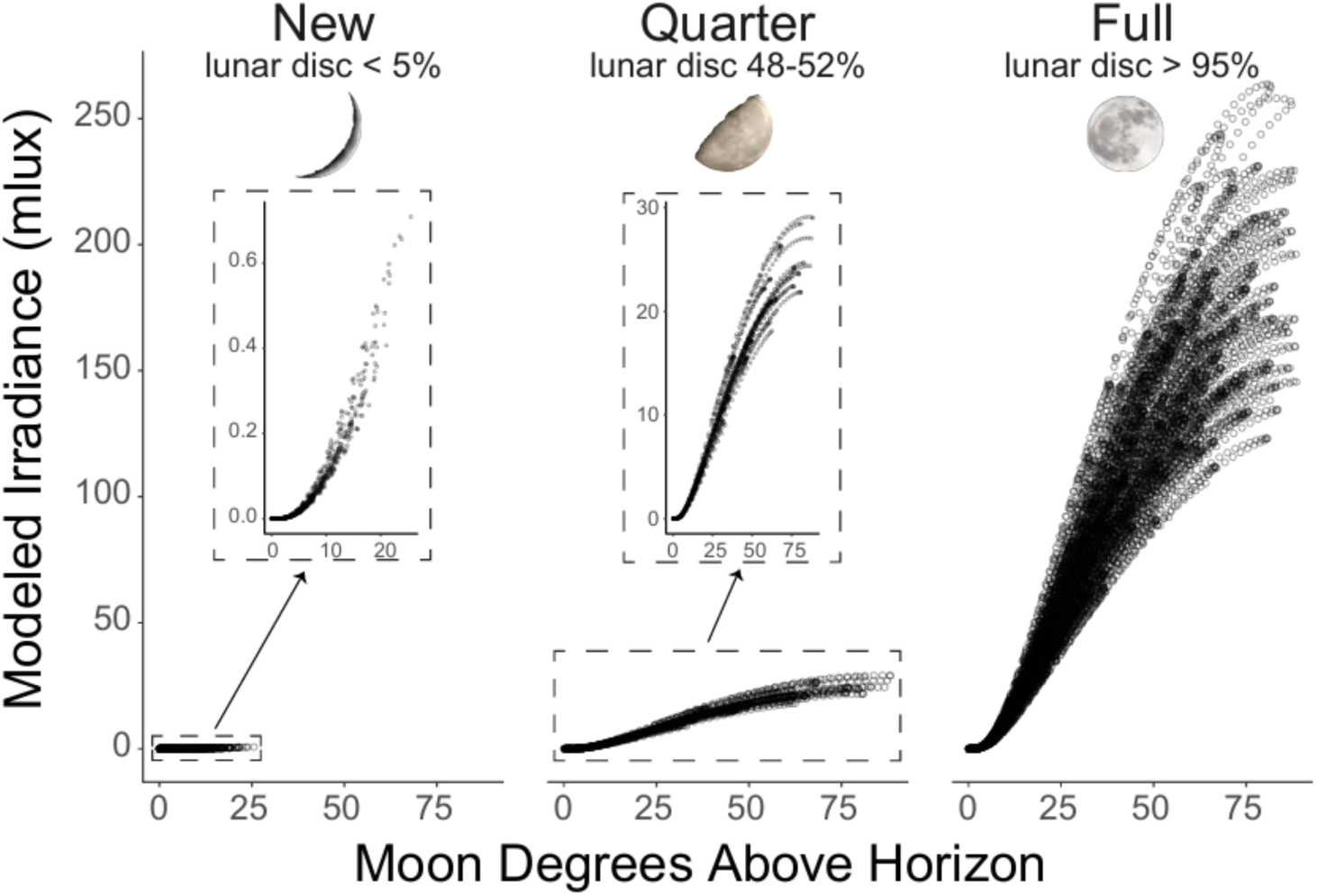
Lunar phase and elevation angle determine irradiance. Panels depict the relationship between lunar elevation angle and modeled irradiance for crescent, quarter, and full moons. Inset graphs in crescent and quarter moon panels contain identical data, but axis ranges are altered to highlight relationship. Note that crescent moons are rarely higher than 10 degrees above the horizon during nighttime.

#### 2.2.3 Spectral components of lunar radiance and irradiance

The spectral composition of moonlight is similar to sunlight, but slightly long-shifted (longer wavelengths are more prevalent) due to reflective properties of the lunar surface (Lane and Irvine 1973, Kieffer and Wildey 1996). Moonlight’s spectral composition does not change significantly across the circa-monthly lunar cycle (Lane and Irvine 1973, Miller and Turner 2009). However, the spectral composition of lunar irradiance varies within the nighttime, depending on the lunar elevation angle above the horizon. Due to increased atmospheric scattering, moonlight is slightly more red-shifted near moonrise and moonset.

#### 2.2.4 Lunar Polarization

The night sky polarization pattern, created by scattering of moonlight in the atmosphere, is analogous to the pattern around the sun in the daytime sky – the primary difference is its weaker intensity (Cronin and Marshall 2011). The highest degree of polarization in the sky is found in a band 90 degrees from the moon, as with the sun (Cronin and Marshall 2011, Foster et al. 2019). The strength of the polarization signal in the night sky varies across the lunar cycle such that the degree of linear polarization increases with the fraction of the moon that is illuminated (Foster et al. 2019). The polarization signal can be weakened by cloud cover, atmospheric turbidity, and artificial light pollution (Kyba et al. 2011b, Foster et al. 2019), which may be particularly relevant when the moon is near the horizon (Kyba et al. 2011b).

### 2.3 Organismal Response to Moonlight

Moonlight is an ecological variable composed of non-mutually exclusive physical values (irradiance, radiance, polarization) that vary across multiple temporal scales (circa-monthly with lunar phase, daily with lunar elevation). These characteristics of moonlight follow predictable cycles and are important drivers of crepuscular and nocturnal ecology (Kronfeld-Schor et al. 2013, Gaston et al. 2017, Seymoure et al. 2025). The best-studied organismal response to lunar conditions is spatiotemporal activity patterns. Lunar modulation of activity has been documented in terrestrial vertebrates including birds (Jetz et al. 2003, Evens et al. 2020, Dickerson et al. 2020, Alonso et al. 2021, Hedenström et al. 2022), mammals (Hecker and Brigham 1999, Saldaña-Vázquez and Munguía-Rosas 2013, Pratas-Santiago et al. 2017, Roeleke et al. 2018, Taylor et al. 2023), non-avian reptiles (Lillywhite and Brischoux 2012), and invertebrates including insects (Bidlingmayer 1964, Kerfoot 1967, Youthed and Moran 1969, Corbet et al. 1974, Lang et al. 2006, Gunn and Gunn 2013, Young et al. 2021), arachnids (Tigar and Osborne 1999, Nørgaard et al. 2006), and decapods (Saigusa 1980). Reproductive behavior has been linked to lunar periodicity in various taxa, including insects (Nemfc 1971), mammals (Peeva et al. 2023), and birds (Mills 1986). Many organisms, particularly arthropods, utilize the moon for orientation and navigation (Foster et al. 2019, Torres et al. 2020, Storms et al. 2022, Freas and Cheng 2025). The variation in moonlight across lunar cycles may exert phenotype-dependent pressures on individuals within species and populations (San-Jose et al. 2019). For additional discussion of the ecological relevance of moonlight, we suggest two excellent review papers (Kronfeld-Schor et al. 2013, Prugh and Golden 2014).

## 3 Methods

In this manuscript, we review published studies on ecological impacts of ALAN on animals. First, we qualitatively review how researchers have incorporated moonlight into these studies. Second, we review and summarize findings on the interactive effects of ALAN and moonlight. Informed by the results of this literature review, we then outline general hypotheses regarding the interaction of moonlight and ALAN.

### 3.1 Literature Search

We performed a keyword literature search using The Web of Science (‘All databases’ option) on papers published before November 2025. We used the terms [(‘artificial light* at night OR light pollution) AND (‘species’ OR ‘ecolog’ OR ‘predat’ OR ‘abundance’ OR ‘behavior’ OR ‘activity’ OR ‘flight to light’ OR ‘pollination’)]. Additionally, we surveyed reference lists of ALAN review papers to incorporate papers not detected by the WOS search. We restricted our focus to animals, excluding studies on plants. We excluded studies on subaqueous ecology. We did not employ a rigid taxonomic demarcation: if a study investigated events occurring terrestrially or aerially, we included it, even if stages of that organism’s life cycle are aquatic. We focus primarily on field-based experimental or observational studies. Though there have been advances in simulating moonlight conditions in laboratory settings (Poon et al. 2024), the majority of lab-based studies lack inclusion of lunar information for practical reasons. We retained 379 journal articles for review. We did not review supplementary materials or review papers for methodological accuracy.

### 3.2 Paper Categorization

We categorized papers by taxonomic focus and lunar inclusion. We classified taxonomic focus as birds, mammals, reptiles, amphibians, arthropods, other invertebrates, and multiple/non-specific. We created a lunar inclusion classification scheme wherein each paper was scored as (1) no lunar inclusion, (2) experimental blocking, (3) categorical inclusion or (4) numeric inclusion. Papers in the first group did not incorporate moonlight into their study design. Papers in the second group sought to reduce confounding effects of moonlight by performing all data collection under similar lunar conditions. In the third group, moonlight was treated as a categorical variable (e.g., data collected during new and full moon). In the fourth group, moonlight was treated as a numerical variable (i.e., lunar elevation angle, lunar disc %, or lunar irradiance). Due to small sample size, the diversity of study systems and scales, and variability in quantification of light environments, we did not perform a quantitative meta-analysis. Rather, for papers that incorporated lunar variables, we recorded if (1) the study tested an ALAN-moonlight interaction and (2) if an interaction was reported.

The precision of lunar information generally increases from group 1 to group 4. Studies blocking for confounding effects of moonlight better account for lunar information than studies without lunar information. Incorporating moonlight as a categorical or numerical variable provides more information than experimental blocking. Numerical inclusion retains greater precision than categorical inclusion. For example, modeling the effect of lunar elevation and lunar irradiance provides more information than a binary of new vs full moon. This classification does not equal a ranking of merit. Each group contains studies with a variety of methodological approaches and research questions. The relevance of moonlight depends on the specific research question; not all studies require detailed, numerical lunar data. There are many challenges in accurately quantifying moonlight in ecological research and the ‘best’ approach may not always be feasible (Kyba et al. 2017).

## 4 Results

### 4.1 Paper scoring

We scored 379 journal articles on taxonomic focus and their inclusion of lunar information in experimental design and analysis (Figure 4A). Arthropods (n = 125), birds (n = 115), and mammals (n = 81) together accounted for 85% of reviewed studies. Insects and bats accounted for 84% (n = 105) and 70% (n = 57) of arthropod and mammal studies, respectively. Reptiles (n = 12) and amphibians (n = 15) together accounted for 7% of studies reviewed. Additionally, 7% of the studies focused on multiple taxonomic groups.

**Figure 4:**
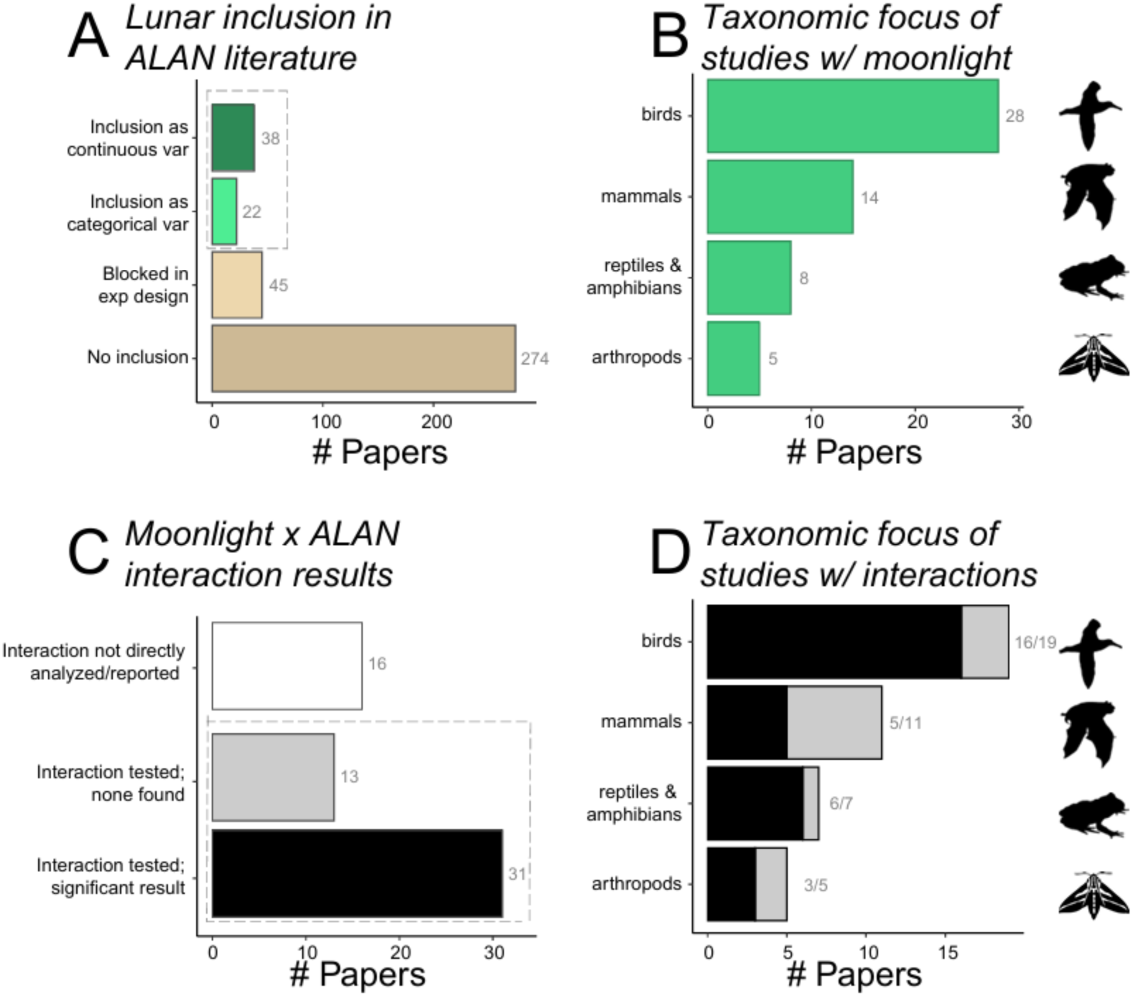
Categorical summary of light pollution literature with respect to lunar inclusion. **A)** Number of papers assigned to each lunar inclusion category. **B)** Representation of major taxon groups in papers that included lunar variables. **C)** Detection of an interaction effect of moonlight and ALAN in papers that included lunar variables. **D)** Representation of major taxon groups in papers that tested interaction of ALAN and moonlight.

Seventy-two percent (n = 274) of these articles did not incorporate lunar data into their experimental design or data analysis. Twelve percent (n = 45) aimed to minimize or eliminate confounding effects of moonlight via experimental design. Sixteen percent (n = 60) incorporated lunar data in experimental design and analysis either categorically (n = 22) or numerically (n = 38).

### 4.2 Lunar inclusion across taxon groups

Lunar inclusion was not consistent across taxonomic groups (Figure 4B). Only 4% of studies on arthropods, but 24% of studies on birds and 17% of studies on mammals incorporated moonlight numerically or categorically. However, 18% of studies on arthropods blocked for confounding effects of moonlight, whereas only 3% of bird studies did the same. Most studies in all main taxonomic groups, except reptiles, did not include lunar data (arthropods, 78%; amphibians, 73%; birds, 72%; mammals, 64%; reptiles, 50%).

### 4.3 Moon x ALAN interaction

Of the 60 papers that included lunar information in their study, 44 explicitly tested an interaction between moonlight and ALAN and 70% (n = 31) reported an effect (Figure 4C). The proportion of papers reporting an interaction was consistently high (Figure 4D) across taxonomic groups (reptiles, 100%; birds, 84%; arthropods, 60%; amphibians, 50%; mammals, 45%).

### 4.4 Trends over time

The number of papers investigating ALAN has increased in recent years (Figure 5A), but the proportion of studies incorporating lunar data has not changed notably. In the past half-decade, the number of papers with no inclusion of lunar data has decreased slightly every year (Figure 5B), but this is largely due to an increase in papers blocking for lunar variables, rather than testing for interactive effects of natural and artificial light.

**Figure 5:**
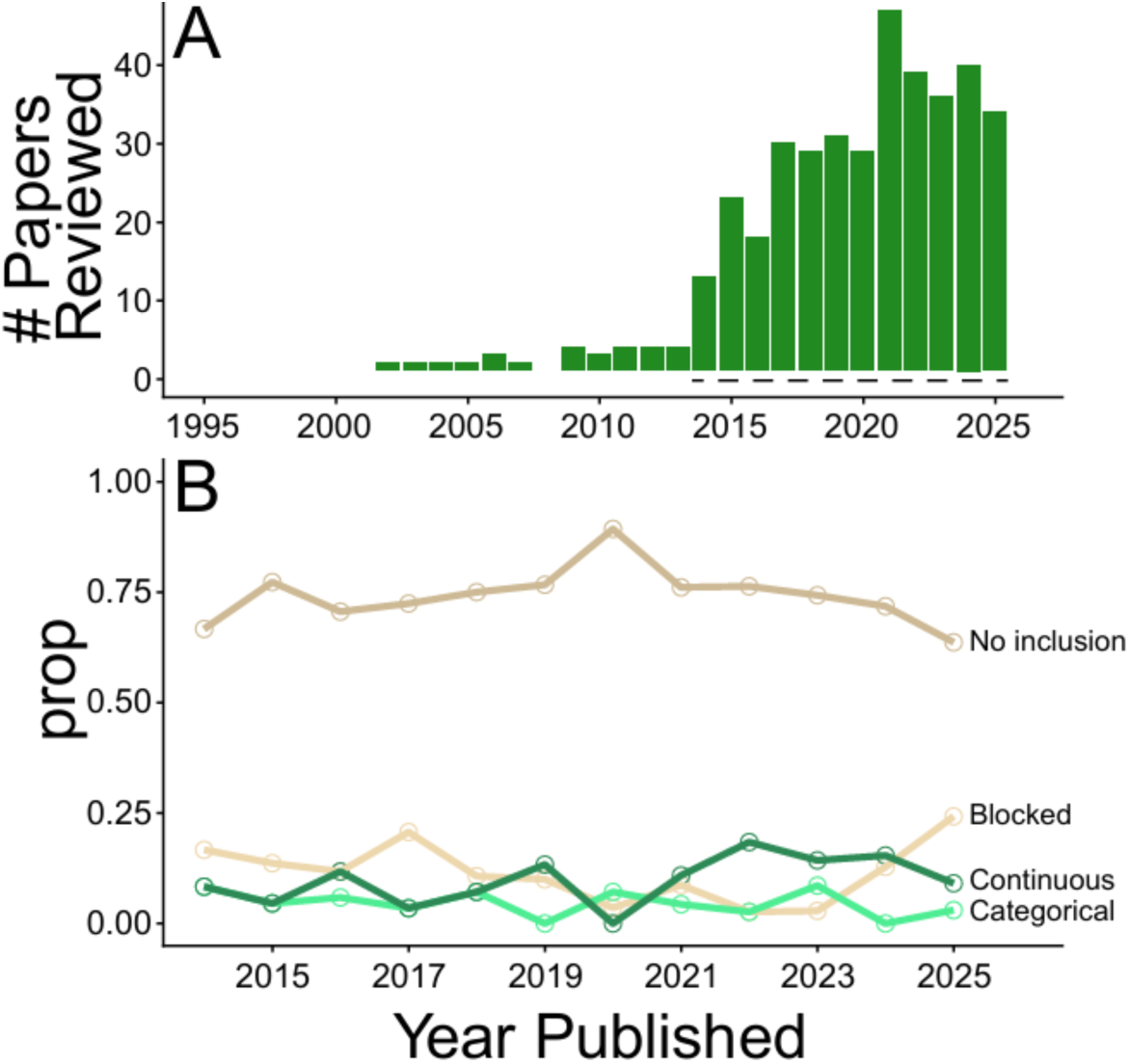
Decadal trend in lunar inclusion in papers reviewed for this manuscript. **A)** Number of papers reviewed by year. Dashed line indicates timespan of panel B. **B)** Proportion of papers per year assigned to each of the four lunar inclusion categories.

## 5 Summary of Literature

### 5.1 Moonlight and ALAN

#### 5.1.1 Disorientation and misorientation

There is consistent evidence across taxa that increasing moonlight reduces disorientation effects from point-source artificial light (Figure 6). Lunar modulation of disorientation has been documented in seabirds (Procellariiformes) (Rodríguez and Rodríguez 2009, Urmston et al. 2022, Rodríguez et al. 2023) and migrating passerines (Passeriformes) (Zhao et al. 2014, Gjerdrum et al. 2021). Disorientation of sea turtle hatchlings (*Caretta caretta*, *Dermochelys coriacea*) by artificial light sources is strongly affected by moonlight, with greater impacts on new moon nights or when the moon is below the horizon (Salmon and Witherington 1995, Bertolotti and Salmon 2005, Berry et al. 2013, Rivas et al. 2015). The inverse relationship between moonlight levels and the number of insects at artificial light sources has been noted for decades (Williams and Singh 1951, Bowden 1981, 1982), but has been primarily contextualized around insect sampling strategy rather than ecological impacts of ALAN (Yela and Holyoak 1997). This well-established relationship appears to have shaped lunar inclusion in ALAN research: a higher percentage of studies on arthropods, relative to other taxonomic groups, block for moonlight while a lower percentage explicitly incorporate lunar data. The ecological significance of the interaction between moonlight and point sources likely extends far beyond raw numbers of insects at artificial lights. For example, one study found that artificial light sources function as a barrier to movement for Lappet moths (Lepidoptera: Lasiocampidae) only when the moon is below the horizon (Degen et al. 2024). Similarly, a study reported that dung beetle (*Scarabaeus satyrus*) orientation behavior was less altered by point-source ALAN on moonlit nights (Foster et al. 2021).

**Figure 6.**
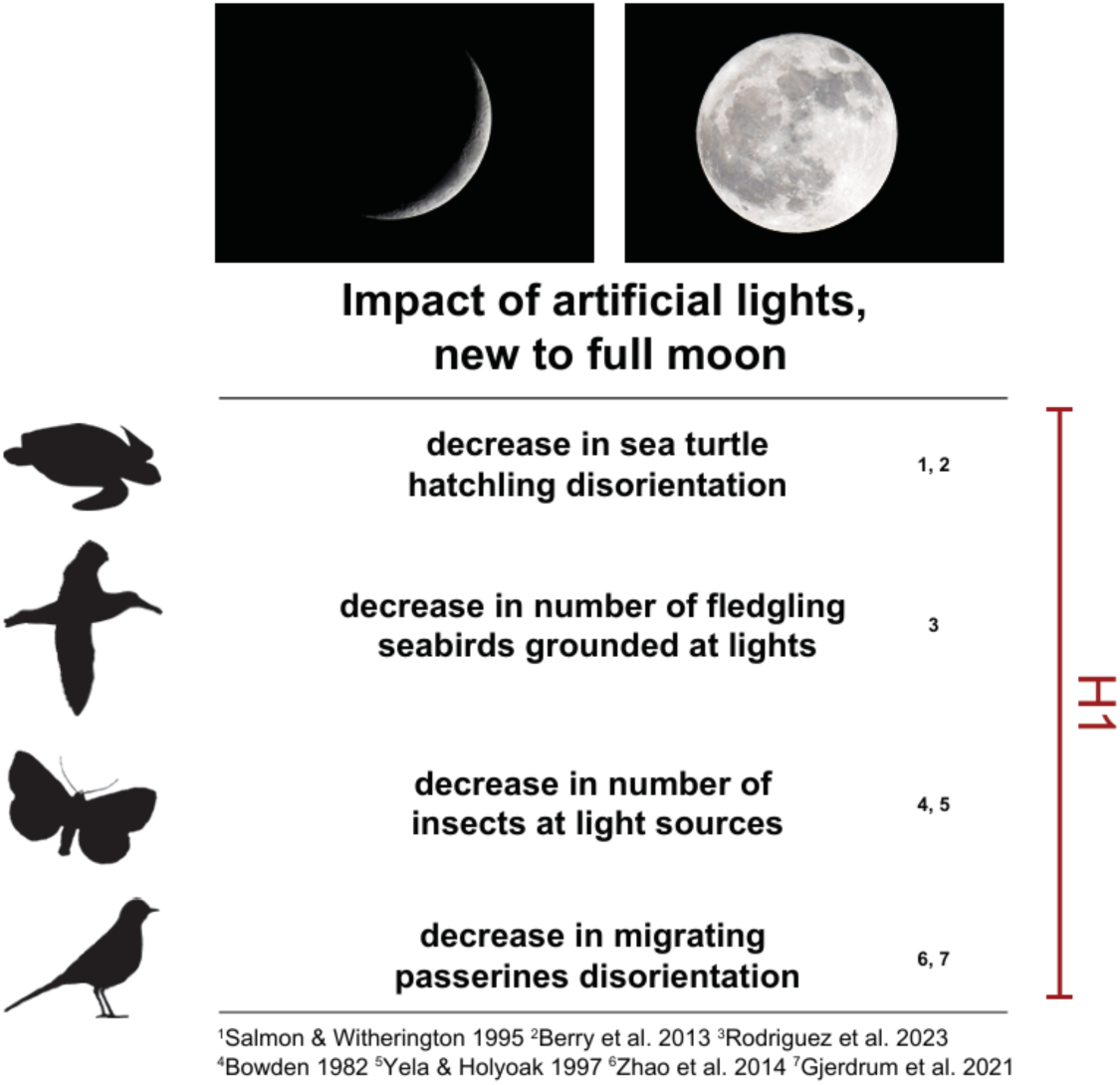
Examples in literature of lunar modulation of the impacts of point source ALAN. Silhouette images downloaded from phylopic.org under the CC0 1.0 Universal Public Domain license.

Skyglow causes disorientation and altered navigational behavior in organisms, though the mechanism may differ from that of point sources (Foster et al. 2021). Organisms utilize various celestial cues in navigation, including bright stars, the milky way, the moon, and the lunar polarization gradients (Foster et al. 2019). Foster et al. 2021 performed a series of experiments untangling the response of dung beetle (*Scarabaeus satyrus*) orientation behavior in response to skyglow, point sources, and celestial cues. Dung beetles responded to skyglow by abandoning celestial cues and orienting by terrestrial cues, including nearby artificial light sources. In the absence of direct light sources, orientation was poor under skyglow, indicating that skyglow is not itself used as a cue. However, on moonlit nights, beetles displayed adequate orientation behavior even in the presence of skyglow.

#### 5.1.2 Activity

Interactive effects of ALAN and moonlight on activity patterns have been documented in a variety of taxonomic groups (Figure 7). Nightjar (*Phalaenoptilus nuttallii*) calling behavior was positively correlated with increasing lunar phase in the absence of ALAN, but this synchronicity was eliminated by ALAN (Preston and Brigham 2023). Similarly, nocturnal singing activity in a primarily diurnal songbird (*Rhipidura leucophrys*) was positively correlated with lunar phase in areas with low skyglow, but negatively correlated in areas with high skyglow (Dickerson et al. 2020). Lunar conditions influence activity in many bat species, lunar phobia being common (Prugh and Golden 2014), but the relationship between moonlight and activity can be weakened or eliminated by ALAN (Mariton et al. 2022, Li et al. 2024). Similarly, one study found that bat activity differed between dark and ALAN sites on nights with a new moon but not on nights with a full moon (Frank et al. 2019). In several rodent taxa, specifically voles (*Myodes glareolus*) and kangaroo rats (*Dipodomys stephensi*), difference in activity between ALAN and dark conditions decreased with increasing moonlight (Hoffmann et al. 2018, Shier et al. 2020). Marten (*Martes martes*, *M. foina*) activity in natural habitats varied with lunar conditions but did not in anthropogenic habitats (Wereszczuk and Zalewski 2023), though this study did not experimentally control for light pollution. Gecko (*Tarentola mauritanica*) activity on buildings increased with increasing moonlight, but geckos were active near light sources regardless of lunar conditions (Martín et al. 2018). Outside of the lunar modulation of flight-to-light behavior in insects, few studies have focused on activity patterns of invertebrates in response to moonlight and ALAN. Littoral snails (*Nucella lapillus*) foraged less under full moon conditions in the absence of ALAN, but foraged more under full moon conditions when high levels of ALAN were present (Tidau et al. 2022). Activity in littoral amphipods (*Talitrus saltator*) depended on the interaction of moonlight, artificial light, and cloud cover (Burke et al. 2024).

**Figure 7:**
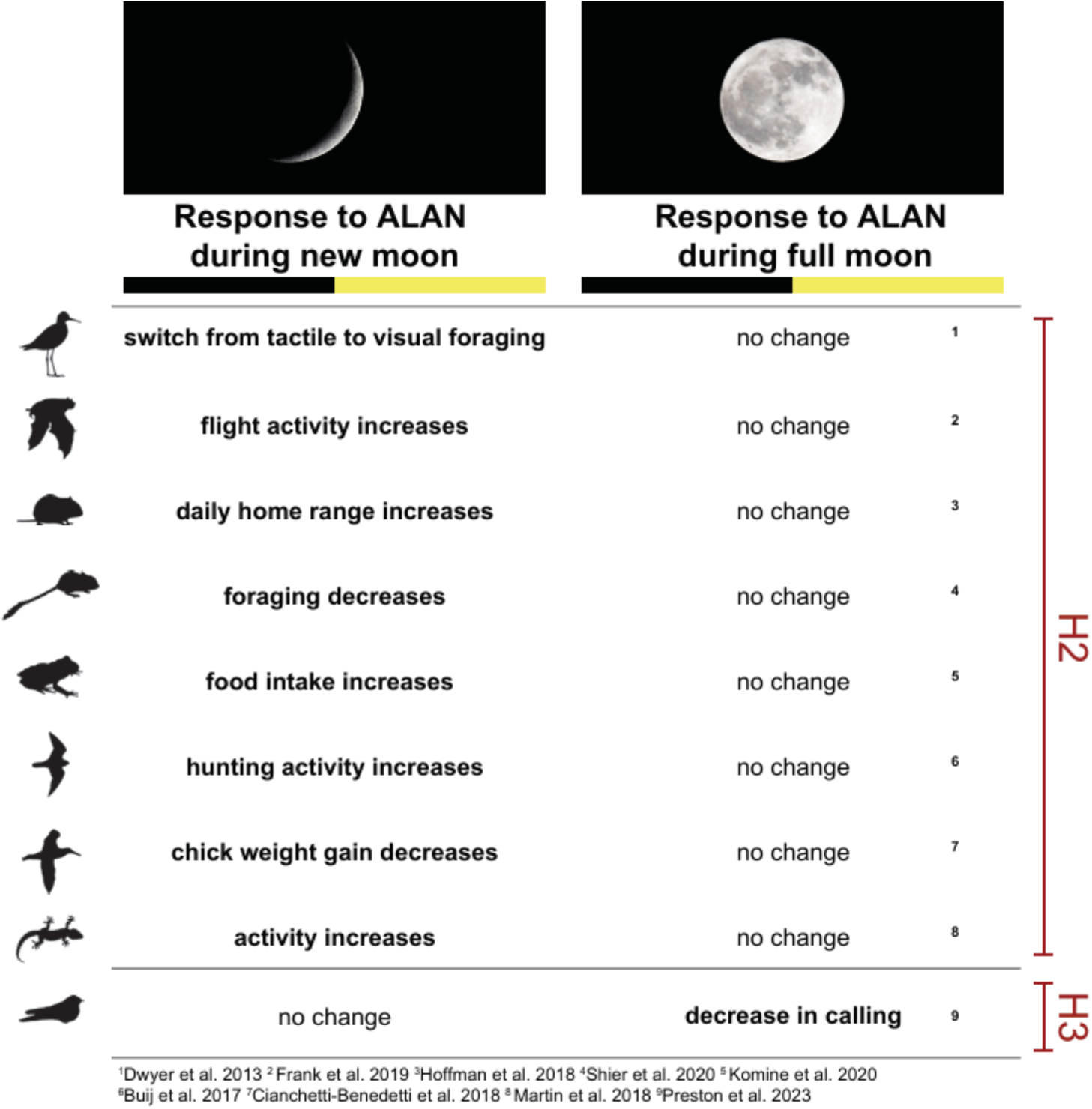
Examples in literature of lunar modulation of the impacts of ALAN as irradiance. Two interaction types: (top) impact of ALAN greater during new moon, (bottom) impact of ALAN greater during full moon. Silhouette images downloaded from phylopic.org under the CC0 1.0 Universal Public Domain license.

#### 5.1.3. Beyond activity and orientation

The interactive effects of moonlight and ALAN extend beyond spatiotemporal activity (Figure 7). Weight gain in shearwater nestlings (*Calonectris diomedea*) was negatively impacted by light and sound disturbances during the new moon, but not full moon (Cianchetti-Benedetti et al. 2018). Redshank sandpipers (*Tringa totanus*) switched from tactile to visual foraging behavior under ALAN when moonlight was minimal but displayed visual foraging with or without ALAN when moonlight was elevated (Dwyer et al. 2013). Anti-predator behavior in curlews (*Numenius arquata*) was influenced by downwelling lunar irradiance only when ALAN irradiance was minimal (Jolkkonen et al. 2023). European nightjars (*Caprimulgus europaeus*) utilized skyglow as a bright visual background for prey detection, but displayed a preference for moonlight if present (Creemers et al. 2025). Cane toad (*Rhinella marina*) foraging efficiency increased near artificial lights, but this effect was reduced by ambient light pollution or moonlight (Komine et al. 2020). A largely-diurnal falcon (*Falco eleonorae*) had greater nocturnal hunting success around artificial lights when moonlight was minimal and conditions were cloudy (Buij and Gschweng 2017). The authors of several studies suggest their results may be linked to lunar modulation of flight-to-light behavior in insects, a source of prey for their respective focal taxa (Buij and Gschweng 2017, Martín et al. 2018, Hoffmann et al. 2018, Frank et al. 2019, Komine et al. 2020).

#### 5.1.4 Studies reporting no interaction

Not all studies reported interactive effects of moonlight and ALAN. Despite the lunar modulation of seabird grounding at artificial light sources, a study reported no differences in body condition of the stranded seabirds (*C. diomedea*) across lunar phases (Rodríguez et al. 2012). One study found that frog (*Lithobates clamitans*) calling behavior was influenced by ALAN, but not by the interaction between ALAN and moonlight (Baker and Richardson 2006). Mountain lions (*Puma concolor*) avoided areas with high upwelling irradiance from ALAN, but no interaction with moonlight was detected (Barrientos et al. 2023). Pinyon mice (*Peromyscus truei*) displayed responses to both moonlight and ALAN, but the interaction between natural and artificial light was not statistically significant (Willems et al. 2021). Several studies on bats failed to find a direct interaction of moonlight and ALAN (Voigt et al. 2018, Luo et al. 2021, Murugavel et al. 2023). The mix of studies detecting and not detecting an interaction suggests that interactive effects of moonlight and ALAN may vary by taxa, the behavior or ecological process under investigation, and additional factors influencing lightscapes.

### 5.2 Other Variables

#### 5.2.1 Cloud Cover

Other than lunar conditions and ALAN footprint, cloud cover is the primary factor determining nighttime lightscapes (Kyba et al. 2011a, Krieg 2021, Burke et al. 2024). Cloud cover differentially influences natural and artificial lighting, typically reducing downwelling irradiance from moonlight and increasing irradiance from ALAN (Kyba et al. 2011a, Krieg 2021). This has direct effects on organismal behavior, though studies incorporating moonlight, cloud cover, and ALAN are scarce. Sleeping behavior in geese (*Branta leucopsis*) changed with cloud cover during new moons but did not change with cloud cover during full moons, perhaps due to cloud cover’s differential effect on natural and artificial light (Van Hasselt et al. 2021). ALAN-induced delay in activity onset in one bat species (*Eptesicus serotinus*) was heightened by cloud cover, but cloud cover had the opposite influence on activity onset in the absence of ALAN (Mariton et al. 2022). Similarly, the response of a crepuscular bird species (*Caprimulgus europaeus*) to cloud cover was reversed in rural and urban areas (Evens et al. 2023).

#### 5.2.2 Biome

Light environments vary across biomes (Cronin et al. 2014, Nilsson and Smolka 2021). Downwelling irradiance is less obstructed by vegetation in open habitats (deserts, grasslands, tundra) relative to closed habitats (forests) (Nilsson et al. 2022). Variation in ground-level irradiance across the lunar cycle may be muted under forest canopies, like the effect of cloud cover and sunlight (Nilsson et al. 2022). Horizontal and upwelling light trespass from artificial point sources is greater in biomes with sparser vegetation (Luginbuhl et al. 2009, Sung 2022). Recently, studies have reported that habitat structure mediates ALAN’s impacts (Eisenbeis et al. 2009, Merckx and Slade 2014, Straka et al. 2019, 2021) and the importance of habitat and moonlight for animal behavior is well documented (Prugh and Golden 2014). The combined interaction of moonlight, ALAN, and habitat remains largely untested. Relatedly, despite advances in modeling propagation of ALAN through heterogeneous landscapes (Aubé 2015), the application of these models to ecological studies is limited, though see (Morrell et al. 2024).

#### 5.2.3 Latitude

Latitudinal variation in the biological impact of light pollution is overlooked (Secondi et al. 2020). Global distribution of ALAN (Secondi et al. 2020) and the activity patterns of animals (Bennie et al. 2014, Wong 2025) vary latitudinally. Lunar cycles are latitudinally synchronous, but important characteristics differ – mean and maximum nighttime lunar elevation, and maximum irradiance, are greater at the equator (Figure 1B,C). The photoperiod is consistent near the equator and seasonally variable at higher latitudes. Consequently, the correlation between lunar phase and other attributes (e.g., nighttime lunar elevation angle) is tighter at the equator (Figure 1A). Climatological conditions are not longitudinally consistent within latitude bands, but there are broad-scale latitudinal patterns in cloud cover (Secondi et al. 2020). For example, the intertropical convergence zone (ITCZ) characteristically has high rates of cloud cover (King et al. 2013).

#### 5.2.4 Season

Light environments are seasonally variable in certain biomes. In deciduous forests, for example, seasonal changes in vegetation structure affect understory light levels – downwelling irradiance is less obstructed in winter due to bare canopies (Song and Ryu 2015, Nilsson et al. 2022). Snow cover increases ambient light levels, reflecting both artificial and natural light (Jechow and Hölker 2019). Changes in vegetation and snow-cover, particularly in urban areas, can lead to seasonal dampening of lunar cycles (Puschnig et al. 2020). Animal ecology changes seasonally due to shifts in behavior or life stage. Many insects, for example, are present as different life stages in different seasons (e.g., eggs and pupae during winter; larvae and adults in summer). Life-stage specific information on ALAN’s impacts is limited for many taxa, but within insects, the impacts of ALAN differ substantially between life stages (Boyes et al. 2020). Some impacts of ALAN are restricted to behaviors such as dispersal of immature organisms that occur in specific seasons (Miller and Rice 2023).

## 6 Hypothesis Framework

The interaction between the lunar cycle and ALAN is not unidirectional. Researchers can start with a relationship between an ecological process and natural light *or* artificial light and subsequently ask how this relationship is modified by the other. In this framework, we largely adopt an ALAN-first approach. Within the lunar-first approach, the story is often unidirectional: the reliability and utility of moonlight as an ecological signal decreases as ALAN increases. However, within an ALAN-first framework, the story is more variable. Additionally, within the time frame of ecological studies, ALAN is a (relatively) constant variable whereas moonlight constantly changes. It is trivial to set ALAN treatments and perform data collection across changing levels of moonlight, but standardizing a fixed level of moonlight in field studies is unreliable due to practical constraints.

Moonlight is a complex ecological variable, but for readability we have simplified to ‘lunar phase’ in the following hypothesis section. The phrase “increasing lunar phase” refers to the progression from new moon to full moon. This increase in lunar phase confers on average greater lunar irradiance, higher lunar elevation angles, stronger night sky polarization signals, and greater proportions of nighttime with a visible moon.

### ALAN as radiance (artificial light sources with physical locations)

The literature consistently reports that impacts of ALAN, encountered as a point of elevated radiance, are modulated by the lunar cycle such that the impact is greater on new moon nights and weaker on full moon nights.

**<u>Hypothesis 1. Light Sources:</u> The impact of ALAN as radiance (artificial light sources) is inversely proportional to lunar phase (Figure 8A). The impact of a light source is dependent on some ratio of light source radiance to ambient light conditions.**

**Figure 8:**
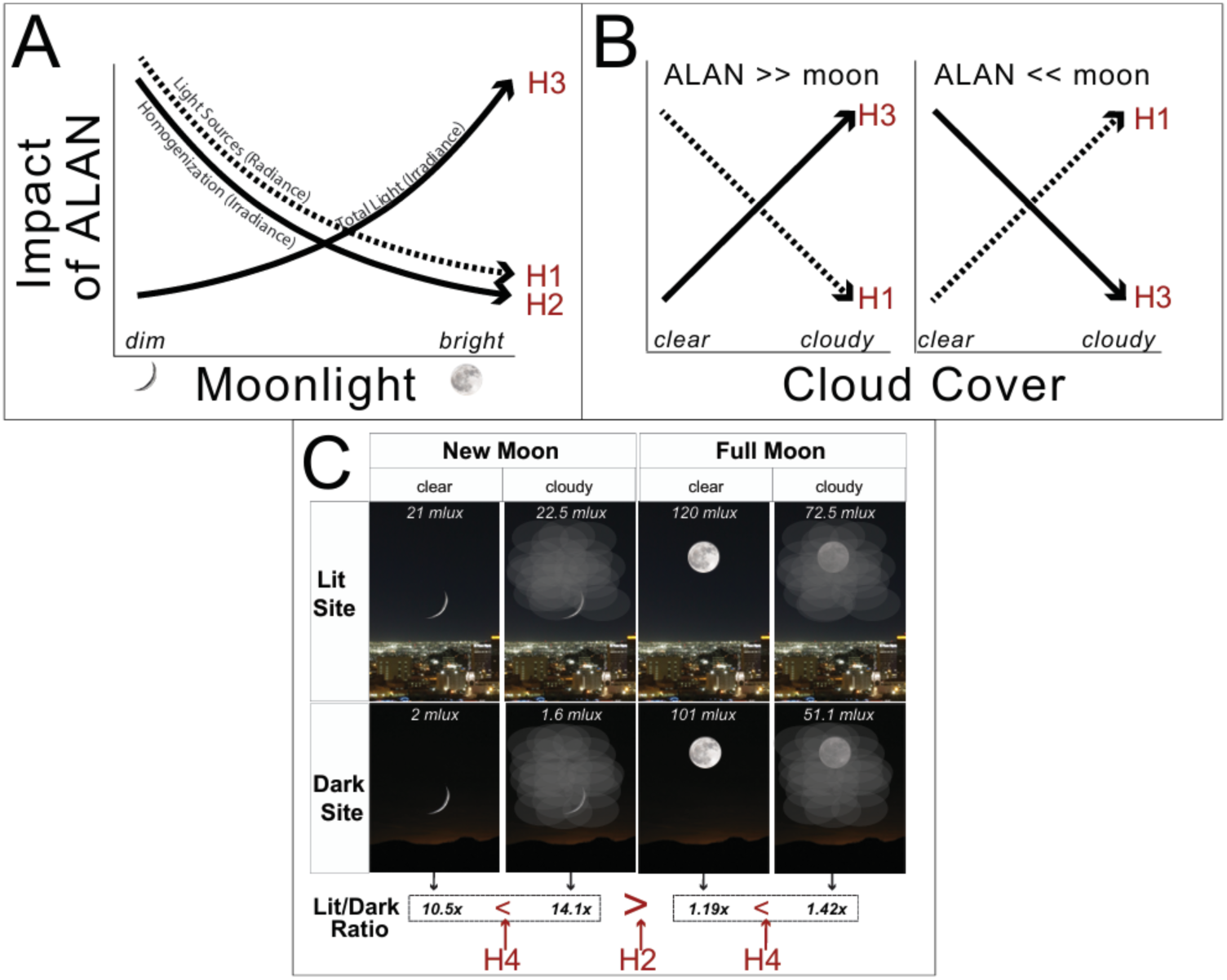
**A)** Three ALAN-moonlight hypotheses (Box 2). **B)** ALAN-moonlight Hypothesis 1 and 3 extended to account for cloud cover. **C)** ALAN-moonlight Hypothesis 2 extended to account for cloud cover. Presented light levels are plausible values chosen to demonstrate interaction of cloud cover, lunar phase, and ALAN at two theoretical sites: a bright site (ambient ALAN of 20 mlux) and a dark site (ambient ALAN of 1 mlux). Full and new moon conditions taken to be 100 and 1 mlux, respectively. Millilux approximated via (1) *total light* = *moonlight + ALAN*, (2) *moonlight with clouds = moonlight * 50%* and (3) *ALAN with clouds = ALAN * 110%*.

The impact of a particular artificial light source seemingly depends on some ratio of light source radiance to environmental light conditions (Bowden 1982, Rodríguez et al. 2023, Battles et al. 2024). The specific feature of a brightened light environment (e.g., background radiance, total downwelling irradiance, scene complexity, sky polarization) that reduces the impact is an open question, and likely varies between taxa (Chan et al. 2025). The suite of lunar characteristics is covariable such that a new moon has reduced radiance, irradiance, and polarization relative to a full moon. Whichever characteristic of the lightscape modulates the impact of ALAN, the ratio between light source and lightscape will decrease as the lunar phase increases. The impact of a light source varies within a given night due to changes in lunar elevation angle, altering lunar irradiance and polarization (Storms et al. 2022). The part of nighttime in which the moon influences the impact of ALAN also shifts throughout the lunar cycle with changes in the timing of moonrise and moonset. Intra-night variation in ecological responses to the interaction of artificial and natural light has received little research attention. Broadly, the impacts of artificial light sources decrease with increasing lunar phase and increasing lunar elevation angle (Bowden 1982, Salmon and Witherington 1995, Yela and Holyoak 1997, Bertolotti and Salmon 2005, Rodríguez and Rodríguez 2009, Rodríguez et al. 2014, 2023, Zhao et al. 2014, Rivas et al. 2015, Gjerdrum et al. 2021, Degen et al. 2024).

### ALAN as irradiance (ambient light level in the environment)

The literature indicates that the impacts of ALAN, encountered as irradiance, are modulated by the lunar cycle. There are two contrasting relationships between lunar phase and the impact of ALAN. Some impacts are greater during new moons (Figure 7, top) and others greater during full moons (Figure 7, bottom). The first relationship was reported by a greater number of studies. We present hypotheses for both relationships. A handful of studies reported more complex responses of organisms to moonlight and ALAN. Intense ambient ALAN can generate perpetual “full moon” conditions. In heavily light-polluted locations, the nocturnal environment is consistently brighter than the brightest natural moonlight conditions, masking circa-monthly and intra-nightly variation in lunar irradiance (Puschnig et al. 2020). Weaker ambient ALAN, too dim to fully mask lunar cycles, still eliminates new moon conditions (Seymoure et al. 2025). The interaction between lunar cycles and ambient ALAN is more complex when ambient ALAN levels are intermediate between new and full moon conditions. We discuss these situations here. For the following hypotheses, we simplify nighttime light levels to the equation *total irradiance* = *moonlight + ALAN*.

**<u>Hypothesis 2. Homogenization:</u> The impact of ALAN as irradiance (ambient light levels) is inversely proportional to lunar phase (Figure 8A). The impact of increased ambient light levels is dependent on the ratio of total light to natural light.**

Under natural conditions, downwelling irradiance varies across several orders of magnitude depending on the lunar phase. ALAN’s contribution to total irradiance homogenizes nighttime light environments by raising light levels to a constant minimum level, independent of moonlight. Most frequently, new and crescent moon conditions disappear; natural variation in light levels is “condensed” towards full moon conditions. Consider two sites (A: 1 mlux of ambient ALAN; B: 20 mlux of ambient ALAN) on two nights (full moon: 100 mlux; new moon: 1 mlux). On the new moon night, site A is 2 mlux and site B is 21 mlux: site B is 10.5 times brighter than site A. On the full moon night, site A is 100 mlux and site B is 120 mlux: site B is 1.19 times brighter. Though the raw difference between dark and lit sites is constant (19 mlux), the percent change from natural light levels is drastically more pronounced on new moon nights. New moon nights are drastically changed by ALAN whereas full moon nights are comparatively less changed. Consequently, we predict that the impact of ALAN is greater on new moon nights or when the moon is low in the sky (Dwyer et al. 2013, Buij and Gschweng 2017, Hoffmann et al. 2018, Frank et al. 2019, Komine et al. 2020, Shier et al. 2020).

**<u>Hypothesis 3. Total Light:</u> The impact of ALAN as irradiance (ambient light levels) is proportional to lunar phase (Figure 8A). The impact of increased ambient light levels is dependent on the sum of natural and artificial light levels.**

The behavior of many animals is dependent on nighttime light levels, exemplified by the importance of moonlight (Kronfeld-Schor et al. 2013, Prugh and Golden 2014) and intensity-specific impacts of ALAN (Miller et al. 2017, Sanders et al. 2018, Dyer et al. 2023). Consider a behavior (e.g., moth oviposition (Nemfc 1971)) that occurs when light levels are below a threshold *L*, and a behavior (e.g., nocturnal singing in a primarily diurnal songbird (La 2012)) that occurs when light levels are above that threshold. As irradiance approaches *L*, p(oviposit) *→* 0 and p(sing) *→* 1. For a given value of ALAN below *L*, irradiance approaches *L* as lunar phase increases. If *L* is greater than maximum natural irradiance due to moonlight, p(oviposit) → 0 and p(sing) → 1 only if ALAN raises total irradiance above *L*. For a given level of ALAN below *L*, *L* is more likely to be reached as lunar phase increases. Within this threshold paradigm, the impact of ALAN as irradiance is predicted to increase with increasing lunar phase (Dickerson et al. 2022, Preston and Brigham 2023). In other words, the impact of ALAN as irradiance is greater on full moon nights and lesser on new moon nights. If artificial irradiance is greater than that of a full moon night, changes in lunar irradiance may not influence the impact of ALAN.

### Skyglow

Skyglow does not neatly fit into one of the three hypotheses above. Skyglow contributes to elevated ambient light levels, and this effect can be dispersed across the landscape due to scattering, causing elevated irradiance far away from the light sources. This aspect of skyglow fits into either Hypothesis 2 or 3. Skyglow may modulate the impact of individual light sources (Hypothesis 1) due to its effect on the entire lightscape. We discuss this aspect within Hypothesis 4, below.

Skyglow is directly measurable as altered radiance or polarization of the sky. Skyglow alters the night sky polarization signal, typically reducing the strength of the degree of linear polarization (Kyba et al. 2011b). Typically, the strength of the lunar polarization signal is weaker on new and crescent moon nights and stronger during gibbous and full moon nights (Foster et al. 2019). Based on these two general patterns, we can hypothesize that skyglow is more likely to lower the nighttime polarization signal below organismal detection thresholds when the signal is naturally weaker. Consequently, the impact of skyglow would be predicted to decrease as lunar phase increases. The interaction between skyglow and the lunar polarization signal depends on the azimuthal direction of the moon and light sources, the elevation angle of the moon, and the height of the sky with skyglow covers. Additional research on organismal response to the interplay of skyglow, lunar cycles, and nighttime polarization is needed.

### Mediation of natural and artificial light by cloud cover

**<u>Hypothesis 4. Cloud Cover:</u> The influence of cloud cover on the impact of ALAN depends on (1) relative intensity of natural and artificial light levels and (2) the form of ALAN.**

Skyglow and cloud cover determine nocturnal light environments (Krieg 2021, Kocifaj and Falchi 2025). Cloud cover typically strengthens ALAN and weakens moonlight (Kyba et al. 2011a, Krieg 2021). If moonlight is the primary contributor to nocturnal light levels at a given place and time, cloud cover will lower light levels. Conversely, if ambient ALAN is the primary contributor of nocturnal light levels, cloud cover will increase light levels. The influence of cloud cover on Hypotheses 1-3 is mediated by the relative amount of ALAN and moonlight in the landscape.

− Hypothesis 1. Light Sources (Figure 8B): Bright nighttime conditions reduce the impact of a particular artificial light source. Though the modulatory effect of skyglow on the impact of point sources is unclear, it has been predicted that higher ambient ALAN may reduce the impact of individual light sources, akin to the effect of moonlight (Wilson et al. 2021, Battles et al. 2024). When ALAN << moonlight, cloud cover will reduce total light levels, thereby increasing the impact of a light source (Yela and Holyoak 1997). When ALAN >> moonlight, cloud cover will increase light levels due to amplification of skyglow, thereby decreasing the impact of a light source (Van Hasselt et al. 2021).
− Hypothesis 2. Homogenization (Figure 8C): We return to our example of Site A (dark) and Site B (ALAN). In clear conditions, Site B is 20 and 1.2 times brighter than Site A on new and full moon nights, respectively. If we consider cloudy conditions that decrease moonlight by 50% and increase ALAN by 10%, Site B is now 45 and 1.44 times brighter than Site A on new and full moon nights, respectively. For both new and full moon conditions, cloud cover increases the difference in light levels between natural and illuminated sites. Consequently, we predict that cloud cover increases these impacts of ALAN irrespective of lunar phase.
− Hypothesis 3. Total Light (Figure 8B): The impact of ambient ALAN is proportional to total light levels. If cloud cover reduces light levels (e.g., moonlit night at a site with low ALAN), the impact of ambient ALAN may decrease. If cloud cover increases light levels (e.g., new moon night at a site with moderate ALAN), the impact of ambient ALAN will increase.

## 7 Mitigation

The interplay between ALAN and the lunar cycle is a critical component of nocturnal ecology. This interaction is further mediated by factors such as weather, habitat, latitude, and season. Understanding these interactions may help mitigate ecological impacts of ALAN. Limiting ALAN during certain phases of the lunar cycle, or specific portions of nighttime, may be an effective mitigation strategy in some systems. This may be particularly effective for spatiotemporally concentrated behaviors, such as sea turtle hatchings, bird migration, and insect mating flights. Lunar-informed lighting decisions is a tool to add to the mitigation toolkit alongside other techniques like light shielding and spectral tuning (Dietenberger et al. 2024, van Koppenhagen et al. 2025, Seymoure et al. 2026).

## 8 Future Directions

There are unanswered questions regarding the interactions of moonlight and ALAN. Topics of interest include (1) scaling of interactions across time, (2) interactions during lunar phases other than full and new, and (3) aspects of moonlight contributing to the modulation of ALAN’s impacts. For some impacts of ALAN (e.g., number of birds grounded at lights), cumulative impact over time intervals can be conceptualized as the integral of discrete timepoints. For other impacts, such as altered species interactions, how do we scale from instantaneous to cumulative impacts? Studies often simplify moonlight to extremely bright (full) or dark (new) nights which represent only a portion of nights. Understanding the importance of moonlight’s modulation of ALAN’s impacts across lunar cycles necessitates quantifying effects across all lunar phases. Lunar phase is a suite of variables like ambient light levels, sky polarization, lunar disc radiance, and height of the moon in the sky. Each aspect is likely relevant to the interaction of the lunar cycle and ALAN. How taxa- or behavior-specific is the importance of different aspects of moonlight? Which aspects are most important to consider in hypothesis generation and experimental design?

## 9 Incorporating moonlight, challenges and recommendations

Studying moonlight ecology in a biologically relevant and optically accurate manner is challenging. Moonlight can be wrangled into a variable for statistical analysis in a plethora of ways and there is a seemingly endless parade of units, terms, and pitfalls when measuring light. Biologically meaningful measures of moonlight are sparse in ecological studies (Śmielak 2023). Many studies report lunar phase, categorically via primary phase terminology or numerically via lunar disc %, rather than field-measured irradiance or radiance values. However, lunar phase is an imprecise proxy for moonlight exposure (Kyba et al. 2020) because lunar phase and irradiance are not linearly proportional; irradiance increases more rapidly with increasing phase near the full moon. Quantifying moonlight categorically groups together nights that have ecologically significant differences in light conditions. This approach also smooths over variation in intra-night timing and duration of moonlight. Ecological studies often use and cite moonlight exposure values that are representative of rare, exceptionally bright full moon nights, unrepresentative of the vast majority of nights (Kyba et al. 2017b).

1. **Consider what aspect of moonlight is relevant for your research objective.** For orientation behavior, important factors to consider might include lunar elevation angle, the lunar polarization signal, and lunar radiance. For temporal activity patterns, the timing of moonrise and moonset may have increased importance. For studies on visual performance (i.e., contrast discrimination, signal detection) the most important value might be lunar irradiance.
2. **We suggest that, for many ecological studies, the two most important lunar values to report are the mean intensity and duration of moonlight.** For studies where data is collected in discrete time intervals (bat passes at a light, distance traveled by an animal, insects sampled in a light-trap), intensity and duration can be combined into a single moonlight exposure variable. For example, “*proportion of time interval with moon above horizon”* multiplied by *“lunar disc %”* generates a single value accounting for both duration and intensity of moonlight (Willems et al. 2021).
3. **Whenever possible, measure nighttime light conditions in the field.** Lunar irradiance can be approximate with software like the *moonlit* R package (Śmielak 2023). However, due to cloud cover, ALAN, and other site-specific factors, modeled values of lunar irradiance will not exactly represent actual conditions. Lunar elevation data can be calculated reliably with software, but topography and vegetation may alter local lunar visibility. When measuring moonlight exposure, consider what horizon perspective is appropriate for the study. Vertical and horizontal measurements of lunar irradiance differ.
4. **If numerical inclusion of moonlight exposure is not practical, or critical, treat moonlight as a categorical variable with caution.** Interpret results within the context of moonlight as a suite of physical and temporal variables. Consider lunar irradiance, timing and duration of visibility, and polarization in concert rather than solely comparing maximum illumination of each night.

## Conclusion

Only 44 (12%) of the 379 revied papers explicitly investigated the interaction of moonlight and ALAN, but 31 (70%) of these reported an interactive effect. The contrast of these percentages, 12% and 70%, suggests many studies on ALAN are lacking important ecological context. Lunar inclusion is not a requirement for research on ALAN to be valuable, but if a larger proportion of future work on ALAN integrates moonlight, we will arrive at a more complete understanding of how organisms respond to both natural and artificial light at night.

## Data Availability

R code and paper categorization file are available via Zenodo at https://doi.org/10.5281/zenodo.20171208

## Supplementary Materials

### Glossary of Terms

Irradiance: the number of photons (or energy) that strikes a small surface over a short time interval. Thus, light originating from many directions contributes to irradiance. Radiometric units are photons/s/cm^2^ or watts/s/cm^2^. The photometric equivalent is illuminance.
Radiance: the number of photons originating from a specific angular direction that strike a small surface over a short time interval. The number of photons (or energy) is presented in steradians (sr; i.e. three-dimensional angle). The radiometric units are photons/s/cm^2^/sr or watts/s/cm^2^/sr. The photometric equivalent is luminance.
Spectrum: the amount of a range of electromagnetic radiation binned by wavelength or frequency (usually wavelength in biological contexts). A spectrum can be described by intensity, hue, and saturation.
Intensity: the amount of photons (or energy) comprising a spectrum. Generally calculated as the integral of the spectrum.
Hue: indicates which wavelengths contribute most to the spectrum. This is most closely associated with colloquial use of ‘color’. The colors blue and red have hues near 450nm and past 600nm, respectively.
Saturation (or chroma): the measure of how many wavelengths contribute to a spectrum. A spectrum with only one wavelength (i.e. a laser) is monochromatic and completely saturated, whereas a spectrum with all wavelengths equally presented (what we would perceive as white, grey, or black) is achromatic and unsaturated.
Polarized light: light with electric field (e-vector) oscillations aligned in a single direction or plane.
Photometric: wavelengths of light are weighted by the human photopic sensitivity curve.
Luminance: the photometric measure of light emitted from a surface constrained by a solid angle. Photometric equivalent of radiance.
Illuminance: the photometric measure of light striking a surface. Photometric equivalent of irradiance.
lunar phase: the apparent shape of the moon as viewed by an observer on earth (qualitative counterpart to lunar disc illumination %)
lunar disc illumination %: the percentage of the lunar disc reflecting sunlight as viewed by an observer on earth (quantitative counterpart to lunar phase)
Scattering of light: the deflection of light rays from a straight path upon colliding with particles, molecules, or irregularities in a medium. For many particles and molecules, scattering is a function of wavelength, resulting in changes to spectral composition of a stream of light.

